# Limits to the cellular control of sequestered cryptophyte prey in the marine ciliate *Mesodinium rubrum*

**DOI:** 10.1101/2020.07.14.202424

**Authors:** Andreas Altenburger, Huimin Cai, Qiye Li, Kirstine Drumm, Miran Kim, Yuanzhen Zhu, Lydia Garcia-Cuetos, Xiaoyu Zhan, Per Juel Hansen, Uwe John, Shuaicheng Li, Nina Lundholm

**Author notes:** shared first authorship (Andreas Altenburger, Huimin Cai, Qiye Li). The authors declare no competing financial interests.

## Abstract

The marine ciliate *Mesodinium rubrum* is famous for its ability to acquire and exploit chloroplasts and other cell organelles from some cryptophyte algal species. We sequenced genomes and transcriptomes of free-swimming *Teleaulax amphioxeia*, as well as well-fed and starved *M. rubrum* in order to understand cellular processes upon sequestration under different prey and light conditions. From its prey, the ciliate acquires the ability to photosynthesize as well as the potential to metabolize several essential compounds including lysine, glycan, and vitamins that elucidate its specific prey dependency. *M. rubrum* does not express photosynthesis related genes itself, but elicits considerable transcriptional control of the acquired cryptophyte organelles. This control is limited as light dependent transcriptional changes found in free-swimming *T. amphioxeia* got lost after sequestration. We found strong transcriptional rewiring of the cryptophyte nucleus upon sequestration, where 35% of the *T. amphioxeia* genes were significantly differentially expressed within well-fed *M. rubrum*. Qualitatively, 68% of all genes expressed within well-fed *M. rubrum* originated from *T. amphioxeia*. Quantitatively, these genes contributed up to 48% to the global transcriptome in well-fed *M. rubrum* and down to 11% in starved *M. rubrum*. This tertiary endosymbiosis system functions for several weeks, when deprived of prey. After this point in time, the ciliate dies if not supplied with fresh prey cells. *M. rubrum* represents one evolutionary way of acquiring photosystems from its algal prey, and might represent a step on the evolutionary way towards a permanent tertiary endosymbiosis.

## Introduction

Endosymbiotic events have enabled eukaryotes to photosynthesize. More than a billion years ago, during a primary endosymbiosis event, a photosynthesizing cyanobacterium was retained by a non-plastidic unicellular eukaryote. Since then, chloroplasts have spread throughout the eukaryotic tree of life by secondary and tertiary endosymbiosis.

*Teleaulax amphioxeia* is an ecologically important, phototrophic marine unicellular eukaryote (protist) with a worldwide distribution (1). It is 8 – 11 μm long and a member of the enigmatic group of cryptophytes, a group that is challenging to place in the evolutionary tree of life (2). Most cryptophytes have permanent chloroplasts, originating from a secondary endosymbiosis event between a red alga and a phylogenetically distinct, non-photosynthetic host (3, 4). Due to this origin, cryptophyte chloroplasts have a complex membrane topology with four membranes that enclose a nucleomorph between the outer two and the inner two membranes (5–7). The nucleomorph is a highly reduced remnant of the endosymbiotic red algal nucleus. Cryptophytes hence possess DNA of different origin: red algal nuclear DNA in the nucleomorph, chloroplast DNA, cryptophyte mitochondrial DNA, and cryptophyte nuclear DNA (8).

Being primary producers, phototrophic cryptophytes are at the base of the marine food web, and grazed upon by heterotrophic and mixotrophic protists alike (9). One of these grazers is *Mesodinium rubrum*, an abundant and ecologically important ciliate. *M. rubrum* is widely distributed in coastal ecosystems and known for causing non-toxic red tides (10–12). Acquisition of phototrophy by retaining a chloroplast that originated from a secondary endosymbiosis event is regarded as a tertiary endosymbiosis (13, 14). *M. rubrum* preys on cryptophytes belonging to the genera *Geminigera, Teleaulax* and *Plagioselmis. M. rubrum* cells keep around 20 chloroplasts from its cryptophyte prey, and usually a single enlarged prey nucleus located close to the nuclei of the ciliate (ciliates have two macronuclei and one micronucleus) (15–17). In order to sustain its maximum growth rate of ~0.5 per day, *M. rubrum* has to ingest ~one cryptophyte per day (18, 19). *M. rubrum* covers typically > 98% of its carbon need via photosynthesis at natural prey concentrations, and can replicate the acquired chloroplasts approximately four times after prey deprivation. Eventually, the chloroplasts are degraded, and *M. rubrum* dies unless new cryptophyte prey cells are ingested (18–21). Thus, this tertiary endosymbiosis between a cryptophyte and *M. rubrum* is not permanent and stable, but species-specific (22, 23).

The regulation of cryptophyte genes within *M. rubrum* has previously been studied using RNA-seq, or Expressed Sequence Tags and microarray approaches (24, 25). These studies found a remarkable cellular and metabolic chimerism between host and prey, and showed that *M. rubrum* not only sequesters the organelle machinery of its prey, but also the anabolic potential of the sequestered organelles (25). Most cryptophyte genes involved in photosynthesis were up-regulated after sequestration of the cryptophyte nucleus and chloroplasts into the ciliate (24). However, previous studies had the challenge to distinguish between transcripts originating from *M. rubrum* and transcripts originating from the prey cryptophytes. We used genomic DNA (gDNA) data from free-swimming *T. amphioxeia* and prey-starved *M. rubrum* to overcome this problem. By screening for *k*-mers shared between gDNA reads and transcripts, we were able to assign transcripts to the right species by sequence signature. Using this approach, we could follow the transcriptional changes upon sequestration for cryptophyte and ciliate genes separately. We investigated changes in the level of *T. amphioxeia* genes expressed before and after ingestion by *M. rubrum* and compared those with starved *M. rubrum* cells that had lost the prey nucleus (Fig. 1). We explored changes in the regulation of the sequestered cryptophyte nuclei in response to changing light and time conditions (night, morning and day) corresponding to darkness, 20 minutes after turning on the light, and full light, and focused for the first time on transcriptional changes of ciliate genes under different light conditions and prey availabilities.

**Fig. 1.**
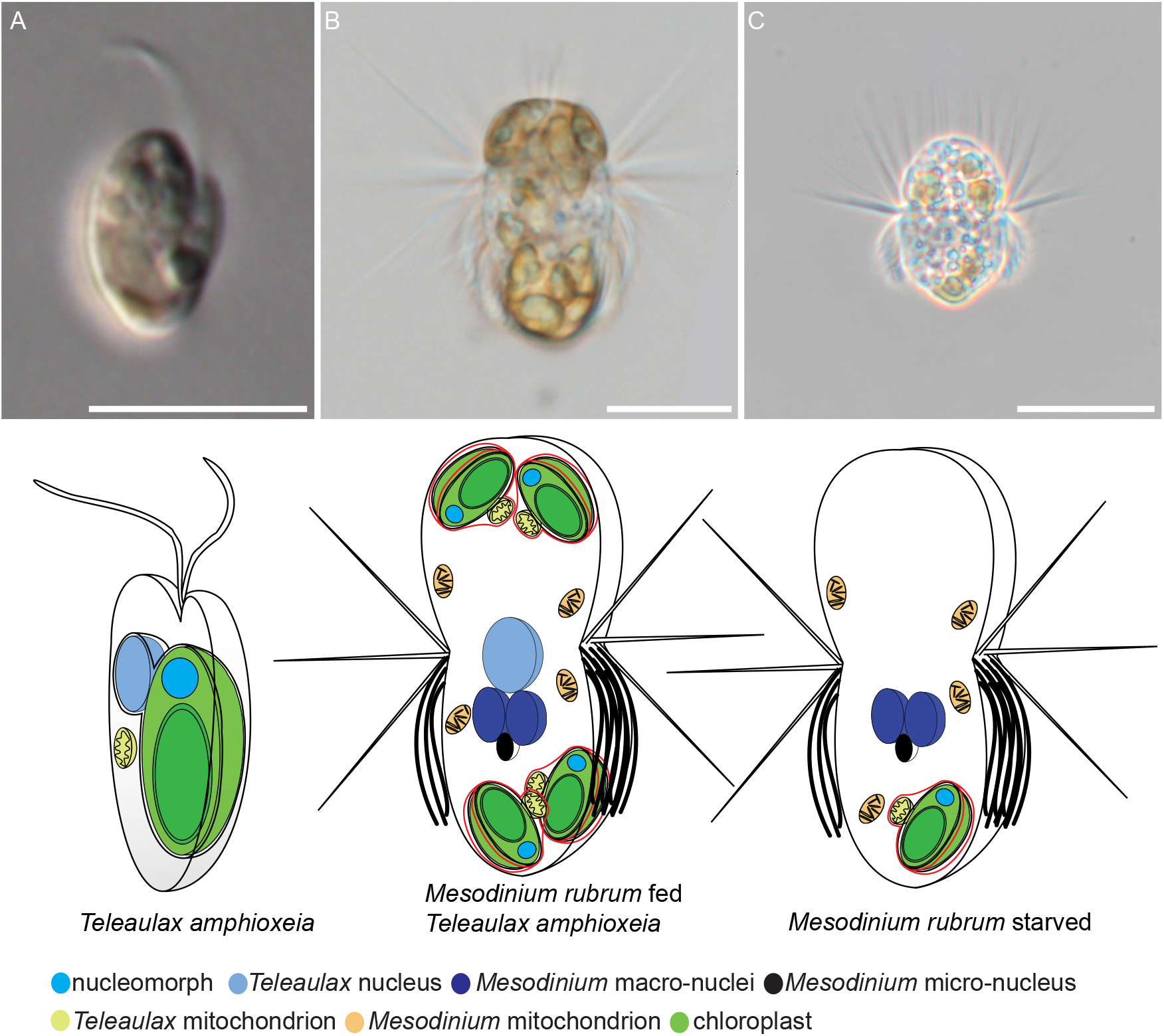
Light micrographs of *Teleaulax amphioxeia* and *Mesodinium rubrum* with corresponding cartoons. (*A*) free swimming *T. amphioxeia* with chloroplast, nucleomorph, mitochondrion and nucleus. The outer membrane of the nucleus is connected to the outer membrane of the chloroplast. (*B*) well-fed *M. rubrum* with two macronuclei, one micronucleus, and one enlarged cryptophyte nucleus. *M. rubrum* contains its own mitochondria, cryptophyte mitochondria, and cryptophyte chloroplasts that are arranged along the periphery of the cell. (*C*) starved *M. rubrum* with two macronuclei, one micronucleus and ciliate mitochondria. Note: starved *M. rubrum* were defined as cultures where at least 90% of cells had lost the cryptophyte nucleus. Note also: Well-fed cells of *M. rubrum* have one enlarged cryptophyte nucleus, which is always located in the center of the cell, termed CPN (centered prey nucleus) (24). Well-fed cells might keep some extra prey nuclei in the periphery of the cell. Upon ciliate cell division, one of the two daughter cells receives the CPN, while in the other, one of the extra prey nuclei migrate close to the ciliate nuclei and enlarges (16). Scale bar equals 5 μm in (*A*), and 10μm in (*B*) and (*C*).

## Results and Discussion

### Transcriptomic profiles and reference gene set constructions for *Teleaulax amphioxeia* and *Mesodinium rubrum*

We performed RNA-seq on cultures of free-swimming *T. amphioxeia*, *M. rubrum* well-fed, and *M. rubrum* prey-starved for more than 4 weeks (i.e. more than 90% of the cells in the *M. rubrum* culture had lost the central cryptophyte nucleus) (Fig. 1). Each culture was sampled at three time points during the light dark cycle: night (6 hours after the light was switched off), morning (20 minutes after the light was switched on), and day (7 hours after the light was switched on) (Fig. 2A). Biological triplicates were collected for each condition, and an average of 184 million reads were generated for each biological replicate (Supplementary Table 1). To accurately discriminate the species-origin of each assembled transcript, we also performed genome sequencing for DNA extracted from free-swimming *T. amphioxeia* and starved *M. rubrum*, respectively (Fig. 2A; Supplementary Table 1).

**Fig 2.**
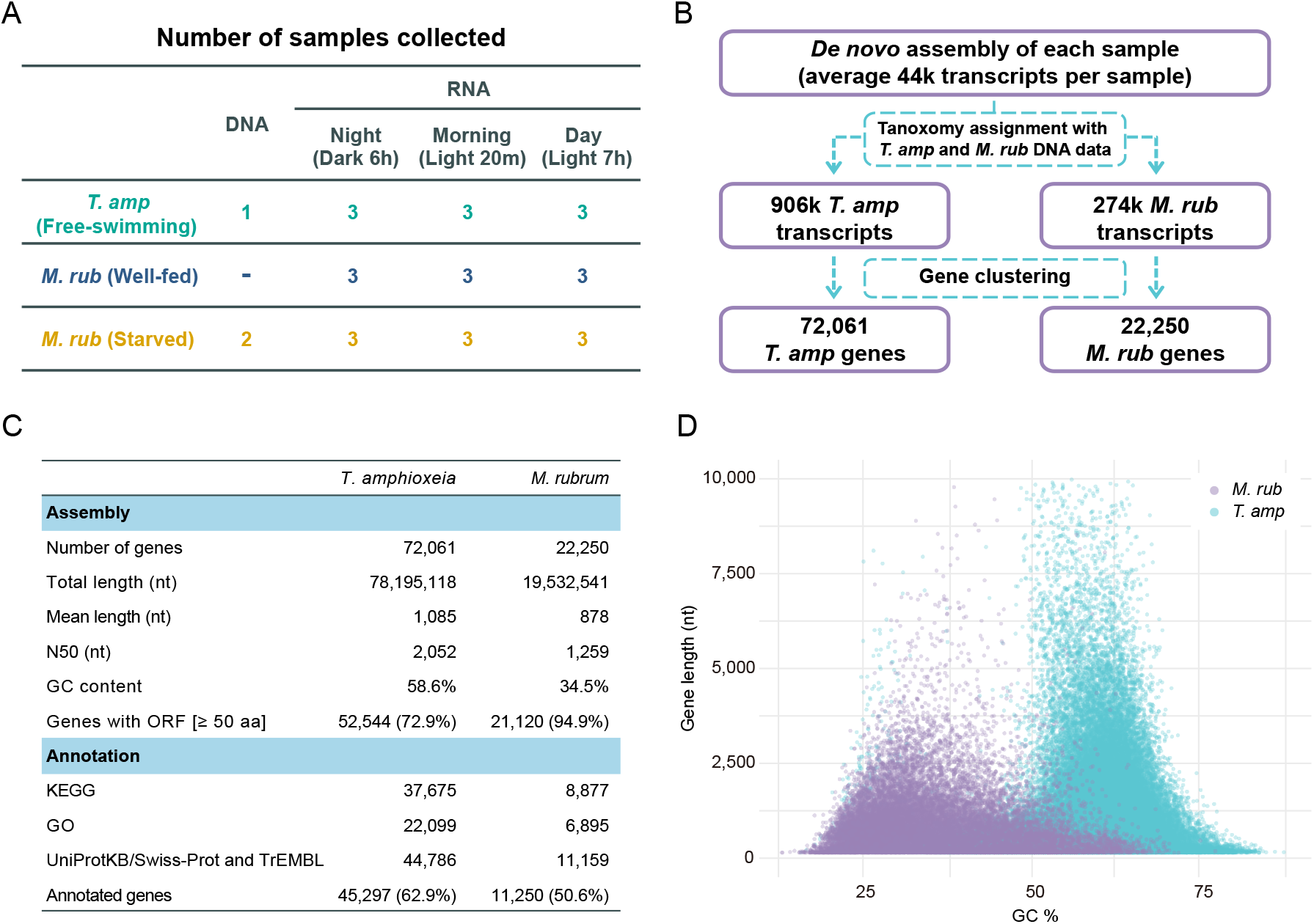
Workflow and transcriptome features of *Teleaulax amphioxeia* and *Mesodinium rubrum*. (*A*) Sampling strategy. (*B*) Analysis workflow. (*C*) Summary of the non-redundant reference gene sets constructed from the *de novo* transcriptome assembly. Abbreviations: nt, nucleotides; aa, amino acids; ORF, open reading frame; KEGG: Kyoto Encyclopedia of Genes and Genomes; GO: Gene Ontology. (*D*). comparison of gene length and GC content for *M. rubrum* and *T. amphioxeia* genes respectively.

As no reference nuclear genomes were available for *T. amphioxeia* and *M. rubrum*, we *de novo* assembled the transcriptome of each sample separately in a first step, followed by stepwise combining the transcripts assembled from each sample. The species identity of each transcript was determined by screening the *k*-mers shared between gDNA reads and transcript sequences (Fig. 2B; Supplementary Table 2; see methods for details). This allowed us to identify 72,061 and 22,250 non-redundant transcripts (i.e. genes) as *T. amphioxeia*- and *M. rubrum*-origin, respectively (Fig. 2C). To access the representativeness of the reference gene sets, we aligned the RNA-seq reads from free-swimming *T. amphioxeia* samples to the 72,061 *T. amphioxeia* genes, and aligned the reads from *M. rubrum* samples to the collection of 72,061 *T. amphioxeia* and 22,250 *M. rubrum* genes (Note: *M. rubrum* samples transcribed genes from both the host and prey genomes). On average, 97.7% of the reads (ranging 96.2% - 98.6%) could be mapped back to the reference gene sets, 95.7% (ranging 93.1% - 97.2%) were aligned in proper pairs, and 88.0% (ranging 84.9% - 89.6%) had mapping quality ≥ 30 (Supplementary Table 3), demonstrating that most sequences in the transcriptomes are present uniquely in the two reference gene sets. We also aligned the RNA-seq reads from free-swimming *T. amphioxeia* samples to the collection of *T. amphioxeia* and *M. rubrum* genes, and observed less than 0.02% of the aligned reads being mistakenly mapped to *M. rubrum* genes, highlighting the reliability of our DNA-based species assignment process.

We annotated 62.9% of *T. amphioxeia* and 50.6% of *M. rubrum* genes by searching against different functional databases (Fig. 2C). Interestingly, the GC content of *T. amphioxeia* genes was around 59%, thus considerably higher than the GC content of *M. rubrum* with 35% (Fig. 2C; Fig. 2D). This GC deviation further supports that the genes were assigned to the right species.

### *M. rubrum* keeps all the genetic material and transcribes most genes from the acquired cryptophyte nuclei

By searching for the *T. amphioxeia* genes in the *M. rubrum* gDNA sequence reads, we retrieved almost all (97.3% - 99.9%) of the 72,061 *T. amphioxeia* genes in the two starved *M. rubrum* DNA samples (Fig. 3A; see methods), suggesting that *M. rubrum* keeps all the genetic material from the acquired cryptophyte nuclei. Next, we examined the transcriptional activity of the *T. amphioxeia* nuclei upon sequestration by *M. rubrum*. Gene expression measurement indicated that on average 63% of the *T. amphioxeia* genes were actively transcribed (TPM ≥ 1) inside *M. rubrum* at some time point during the sampling cycle, which comprised 82% of the *T. amphioxeia* genes, when considering all sampling points together (Fig. 3B). Even though these ratios were lower than those observed in the free-swimming *T. amphioxeia* samples (77% in average and 96% in combination), they did indicate that the majority of *T. amphioxeia* genes (82%) were actively transcribed inside the *M. rubrum* cells (Fig. 3B). At the same time, an average of 90% of the *M. rubrum* genes were actively transcribed regardless of light and prey availability (Supplementary Fig. S1).

**Fig. 3.**
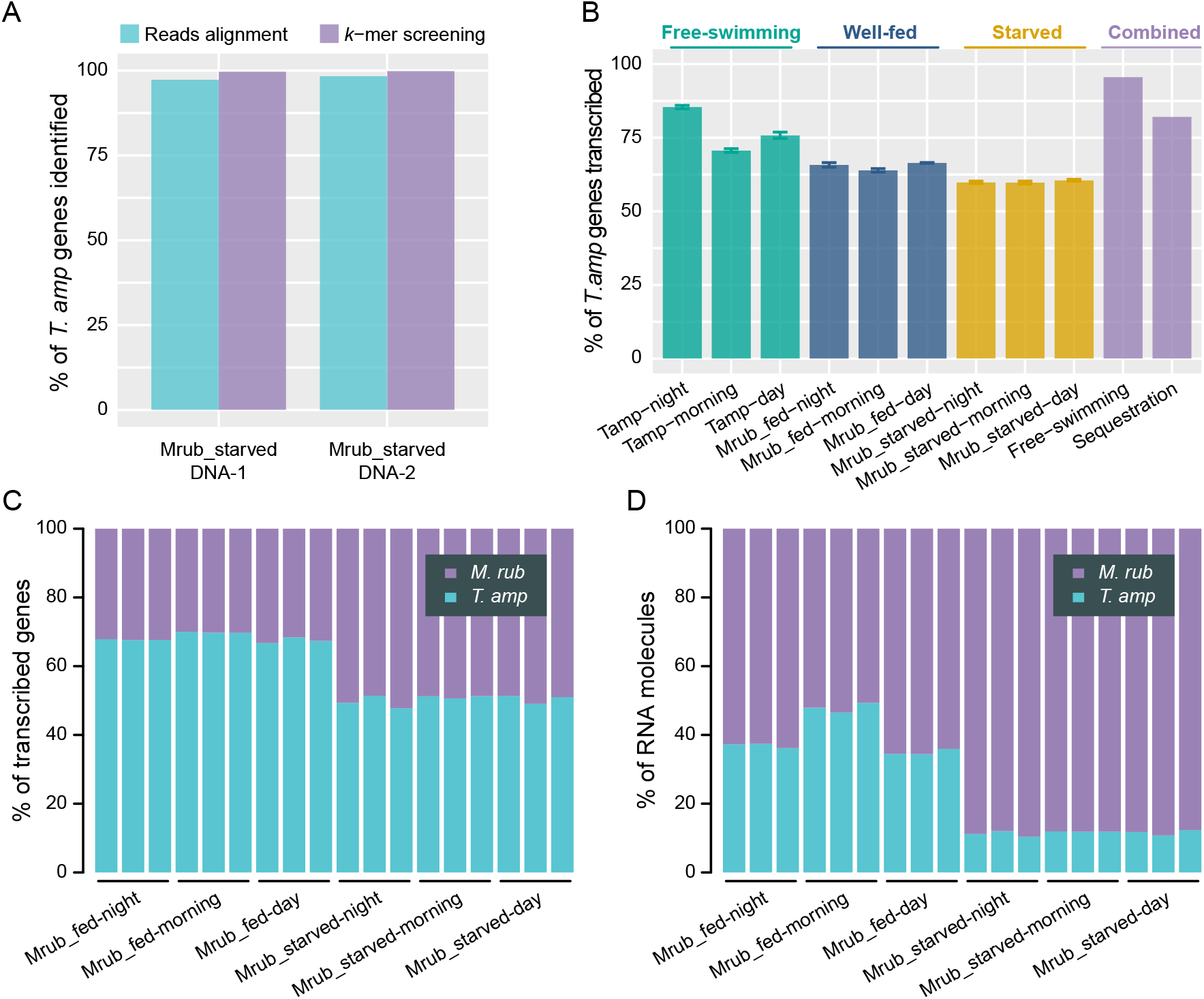
Global transcriptome features of *M. rubrum* (*A*) percentage of *T. amphioxeia* genes identified in the starved *M. rubrum* DNA data by read alignment and *k*-mer screening methods. (*B*) proportion of actively transcribed *T. amphioxeia* genes before and after sequestration. (*C*) global transcriptome of *M. rubrum* with proportion of contributing *T. amphioxeia* and *M. rubrum* genes. (*D*) global transcriptome of *M. rubrum* with proportion of transcript abundance originating from *T. amphioxeia* or *M. rubrum*.

Up to 68.4 ± 1.2% of the genes transcribed within the well-fed *M. rubrum* cells originated from *T. amphioxeia*. This proportion was maintained at 50.4 ± 1.3% for the starved *M. rubrum* samples (Fig. 3C). The contribution of cryptophyte genes to the global transcriptome of well-fed *M. rubrum* in the present study (68.4 ± 1.2%), is higher than previous estimates (13.5% in (24), 58-62% in (25)). However, when taking the transcriptional abundance of each gene into account, the contribution of *T. amphioxeia* transcripts to the global *M. rubrum* transcriptomes was much lower, ranging from 47.5 ± 1.3% (well fed morning) to 10.2 ± 0.9% (starved night) (Fig. 3D). Thus, despite the fact that most *T. amphioxeia* genes were transcribed inside *M. rubrum*, the gene products from *M. rubrum* dominated the mRNA pools of the host cells even in well-fed cells. In a well-integrated endosymbiotic system, one would expect to find a lower qualitative expression of endosymbiont genes: only genes that are beneficial to the host will be expressed, while genes not needed by the host will suffer depletion. Given that 82% of all *T. amphioxeia* genes were expressed at some time point within *M. rubrum*, it is likely that many of the *T. amphioxeia* transcripts are not photosynthesis related and rather by-products. Their functional benefit for *M. rubrum* is not obvious.

### Cryptophyte nuclei present dramatic transcriptional rewiring upon sequestration

Principal component analysis (PCA) with the *T. amphioxeia* gene expression matrix separated all the 27 samples into three distinct clusters of free-swimming *T. amphioxeia*, well-fed *M. rubrum* and prey-starved *M. rubrum* (Fig. 4A), with the distance separating free-swimming *T. amphioxeia* samples from all *M. rubrum* samples being larger than the distance separating well-fed and starved *M. rubrum* groups. This suggests that the condition of sequestration alone induced an overwhelming amount of transcriptional changes when compared with other experimental conditions for the *T. amphioxeia* genes. Consistently, this was supported also by weighted gene correlation network analyses (WGCNA) using the same matrix after filtration for lowly expressed genes (44,241 *T. amphioxeia* genes with mean normalized count ≥ 10) identified six modules. The first two modules comprised 81% of the input genes and enriched *T. amphioxeia* genes that were prevailingly down- and up-regulated after sequestration by *M. rubrum*, respectively (Supplementary Fig. S2). More specifically, differential gene expression analyses between the three sample groups revealed by PCA showed that 34.8% and 31.9% of *T. amphioxeia* genes were significantly differentially expressed (|log_2_FC| > 1.5 and FDR < 0.01) upon sequestration in well-fed and prey-starved *M. rubrum*, respectively (Fig. 4B). All these results consistently emphasize that a profound transcriptional rewiring occurs in the *T. amphioxeia* nuclei after sequestration by *M. rubrum*, as reported previously (24, 25).

**Fig. 4.**
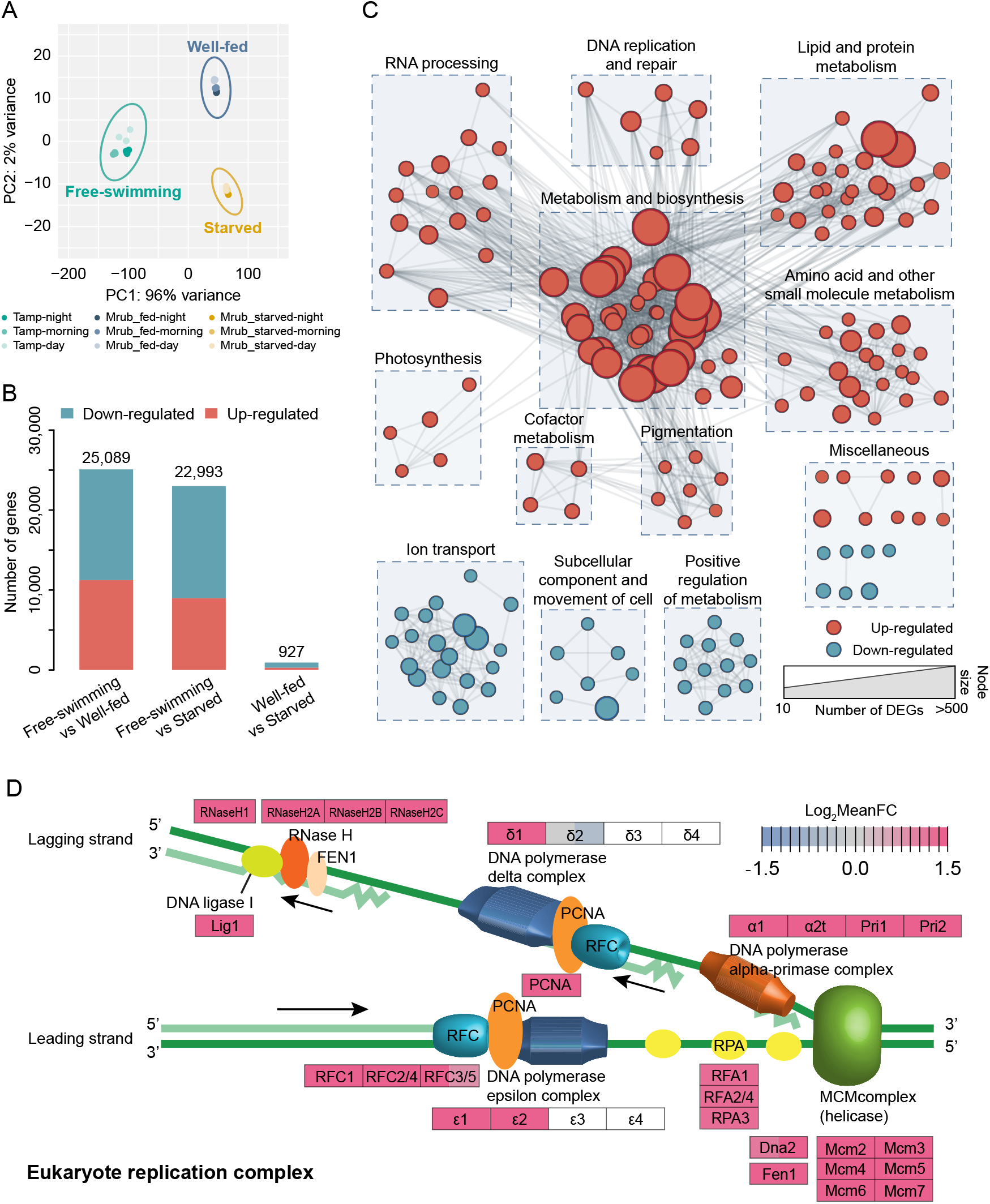
Changes in gene expression of *Teleaulax amphioxeia* genes in response to sequestration. (*A*) principal component analysis of *T. amphioxeia* genes show a clear segregation between free-swimming, well-fed, and starved samples. (*B*) amount of significantly differentially expressed genes (|log_2_FC| > 1.5 and FDR < 0.01) upon sequestration. (*C*) GO enrichment results for *T. amphioxeia* genes up-/down-regulated after sequestration in well-fed samples visualized as an enrichment map. Nodes represent enriched gene-sets and edges represent mutual overlap between gene-sets, thus clustering highly redundant gene-sets. (*D*) changes in *T. amphioxeia* gene expression in the eukaryote replication complex pathway. Left part of each box shows log_2_ fold change in gene expression for free-swimming vs. well-fed samples. Right part of each box shows log_2_ fold change in gene expression for free swimming vs. starved samples.

Functional enrichment analyses for the sequestration-induced differentially expressed genes (DEGs) revealed that *T. amphioxeia* genes related to ion transmembrane transport, signal transduction, cell motility, and regulation of metabolic processes were down-regulated. On the other hand, up-regulation after sequestration was observed for genes involved in photosynthesis, RNA processing, DNA replication and repair, lipid and protein metabolism, and metabolism of diverse compounds (e.g. nucleic acid, carbohydrate, amino acid, carboxylic acid and pigment) (Fig. 4C, Supplementary Table 6, Supplementary Table 7). These results were generally consistent with previous observations by Kim *et al.* (24) and Lasek-Nesselquist *et al.* (25). Interestingly, we also found that the up-regulated DEGs were enriched in DNA replication, repair and recombination, and cell cycle. These comprised *T. amphioxeia* genes encoding cyclins (e.g. CycA, CycH), cyclin-dependent kinases (e.g. CDK2, CDK7), cell division control proteins (e.g. Cdc6, Cdc7, Cdc45) and almost all the genes involved in the eukaryotic replication complex (Fig. 4D, Supplementary Fig. S3). This suggests that the sequestered nuclei are able to replicate their DNA. Besides, in contrast to Lasek-Nesselquist *et al.* (25), we did not observe downregulation of genes involved in protein processing pathways (Supplementary Fig. S4). On the contrary, many genes involved in endoplasmic reticulum membrane and mRNA surveillance pathway were up-regulated after sequestration (Supplementary Fig. S4 and S5). This implies that the sequestered prey nuclei play an active regulatory role in transcription, translation and also in transportation of *T. amphioxeia* gene products.

Of note, only few *T. amphioxeia* genes (927) were identified as DEGs between well-fed and prey-starved *M. rubrum* samples (Fig. 4B), and the majority of DEGs were shared between free-swimming-vs-inside well-fed *M. rubrum* and free-swimming-vs-inside starved *M. rubrum* (supplementary Fig. S6). This demonstrates that the global transcriptional patterns of the *T. amphioxeia* nuclei inside well-fed *M. rubrum* cells were highly similar with those inside starved *M. rubrum* cells.

These findings are unexpected as most of the starved cells had lost their prey nuclei; less than 10% of the starved *M. rubrum* cells had preserved the enlarged centered prey nucleus (CPN). A typical well-fed *M. rubrum* cell has about 20 chloroplasts and a single enlarged CPN, that is located at more or less the same position anterior to the two macronuclei within *M. rubrum* (16). With each chloroplast, *M. rubrum* takes up one cryptophyte nucleus. Well-fed *M. rubrum* cells can contain multiple prey nuclei, i.e. the CPN and some extra prey nuclei that are kept in the periphery of the cell (16). The finding that well-fed and starved cells have similar gene expression patterns, suggests that only the CPN is actively transcribed inside *M. rubrum*. Otherwise, the multiple periphery nuclei have to perform a somehow concerted gene transcription with the CPN inside well-fed *M. rubrum* cells.

Experimental evidence suggests that chloroplasts can divide within *M. rubrum* without the presence of cryptophyte nuclei (16, 18). It is also known that photosynthesis in *M. rubrum* is related to the percentage of cells with a CPN, not to the number of chloroplasts (16). Starved *M. rubrum* cells that have lost the CPN, will usually have some chloroplasts remaining in the cell. A reason why those chloroplasts can survive within *M. rubrum* might be that they are particularly robust, with a comparatively large gene set (26, 27). Likely, the nucleomorph plays a crucial role in enabling the *T. amphioxeia* chloroplasts to divide within *M. rubrum* and renders it a favored prey in comparison to chloroplasts with smaller gene sets (28).

### Cryptophyte responses to light- and time-changes get lost upon sequestration by *M. rubrum*

Free-swimming *T. amphioxeia* is expected to adjust its gene expression pattern according to light and time changes during night, morning and day as any other photosynthetic organism with permanent chloroplasts. This was confirmed by the PCA result (Fig. 4A). To get a closer look at the light- and time-dependent transcriptional responses of the cryptophyte genes before and after sequestration, we conducted pairwise correlation analyses of the free-swimming *T. amphioxeia* samples and the *M. rubrum* samples, respectively. While relatively low correlations were observed among *T. amphioxeia* samples from different light and time conditions (Fig. 5A), we found all the *M. rubrum* samples showing consistently high pairwise correlations (Fig. 5B), implying that the cryptophyte nuclei had lost the ability to adjust gene expression according to light and time changes upon sequestration. This conclusion is further confirmed by the overwhelming amount of dark/light-responding DEGs (10,828; |log_2_FC| > 1.5 and FDR < 0.01) identified in free-swimming *T. amphioxeia* samples in comparison to those identified in *M. rubrum* samples (157; Fig. 5C).

**Fig. 5.**
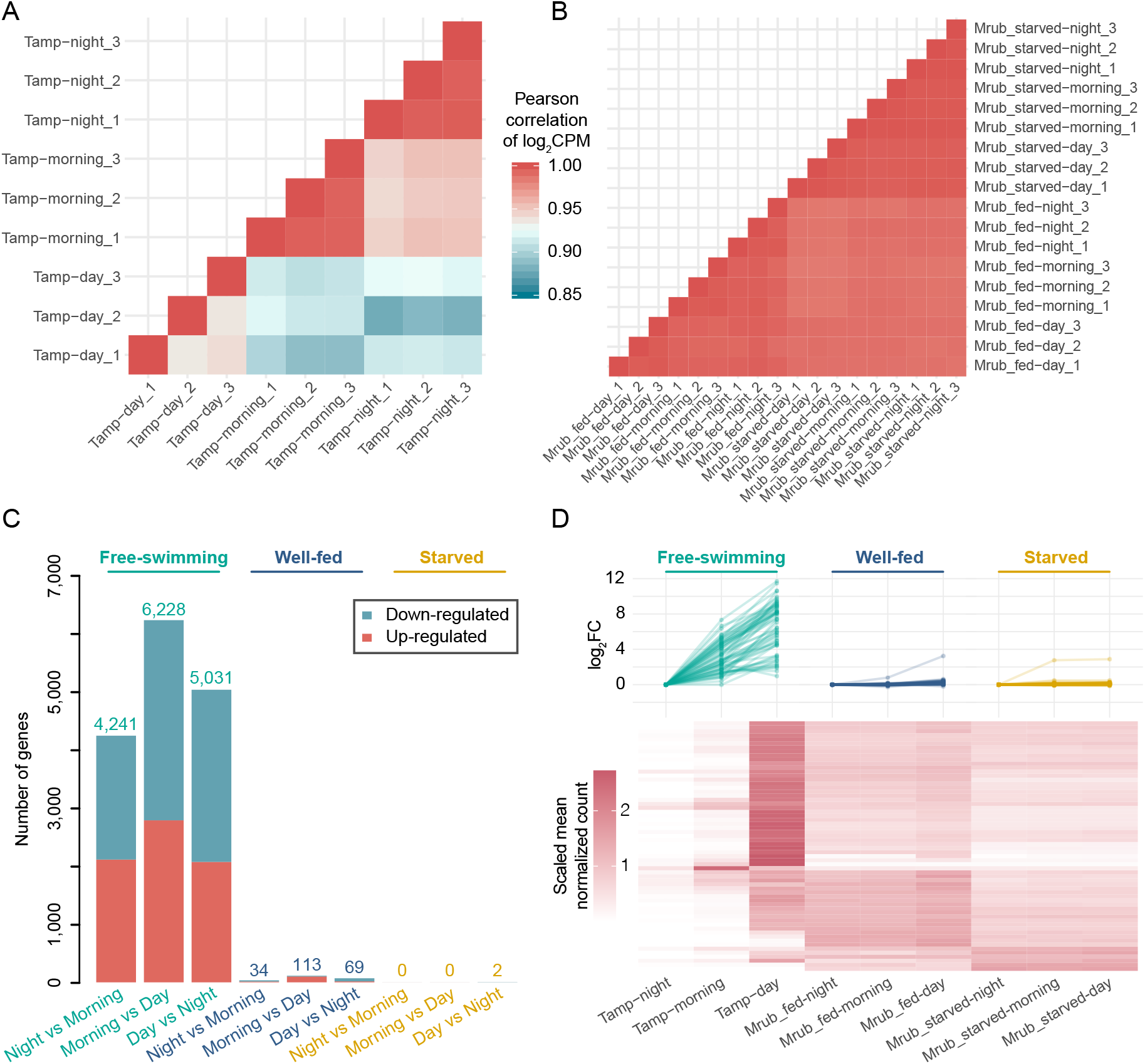
Changes in light and time controlled gene expression of free-swimming *T. amphioxeia* and after sequestration by *M. rubrum*. (*A*) Pearson correlation analysis of *T. amphioxeia* genes among different samples show differences according to time and light condition. (*B*) Pearson correlation analysis of *T. amphioxeia* genes after sequestration by *M. rubrum* reveals an expression pattern that is independent of light and prey availability. (*C*) amount of *T. amphioxeia* genes that were differentially expressed according to time and light condition in free-swimming cells and after sequestration by *M. rubrum.* (*D*) small panels show expression fold change of light dependent *T. amphioxeia* DEGs at night, night-versus-morning, and night-versus-day in free-swimming, inside well-fed *M. rubrum* and inside starved *M. rubrum* condition. The heat map shows *T. amphioxeia* genes that got differentially expressed according to time and light condition before sequestration by *M. rubrum* but maintained at high expression levels at night after sequestration.

*T. amphioxeia* genes, significant differentially expressed according to light and time changes, were functionally enriched in photosynthesis, oxidative phosphorylation, glycolysis and circadian entrainment related pathways (Supplementary Table 8), consistent with the expectation for a free-living photosynthetic organism. Of note, *T. amphioxeia* genes involved in circadian entrainment were generally down-regulated upon sequestration (Supplementary Table 7). This might partly account for the loss of time/light response of the cryptophyte nuclei upon sequestration. The expression of many genes in free-swimming *T. amphioxeia* responded to light (i.e. DEGs up-regulated in morning and day versus night). After sequestration, these light responding genes maintained high expression levels at night. Including DEGs encoding for light-harvesting complex and light reaction of photosynthesis, which are responsible for harvesting and transferring light energy and obviously not needed at night (Fig. 5D, Supplementary Table 9). The loss of the dark/light response together with the over expression of potentially undesired genes strongly suggests that *M. rubrum* can elicit only one expression pattern out of its acquired cryptophyte nucleus regardless of light condition and prey availability (i.e. the number of acquired prey nuclei).

Interestingly, in a different system, the Antarctic Ross Sea dinoflagellate acquires transient chloroplasts from haptophyte prey, and the expression of kleptoplast-targeted genes is also unaffected by environmental parameters such as light (29).

During evolution, foreign chloroplasts have ended up in other protists in many different ways (13). In some protists, intact endosymbionts are well integrated into host cells (30). Other protists reduce ingested algal cells, and keep prey nuclei as well as other cell organelles beside the chloroplasts (like *M.* rubrum). Yet, other protists retain exclusively the chloroplasts for shorter, (i.e. the ciliate *Strombidium*) or longer time (i.e. the dinoflagellate *Dinophysis*) (31, 32). This can be interpreted as evolutionary steps towards permanent endosymbiosis. In a first step, a prey cell is taken up by a host and not digested. In a second step, the host gets some control over the gene expression of the acquired cell via the ingested prey nuclei – that is where *M. rubrum* is right now. In a third step, only the chloroplasts are retained, but need to be replaced with time (ie. *Dinophysis* (33)). In the final step, the genes from the host and the acquired cell (or organelles such as chloroplasts and nuclei) need to align in order to fine-tune the gene expression according to environmental conditions. Whether or not *M. rubrum* is on its way towards a permanent tertiary endosymbiosis is speculative. Such a step will depend on the ability of *M. rubrum* to divide and keep the sequestered prey nuclei permanently, or rely on gene transfer from the algal prey to the ciliate nuclei.

### *M. rubrum* fine-tunes gene expression in response to prey availability and up-regulates genes involved in transport when well fed

The problem that *M. rubrum* faces is that it has to deal with different genetic codes. Ciliates show deviations in the genetic code and it has been suggested that these deviations have occurred multiple times independently (34). *M. rubrum* uses a genetic code that is different from cryptophytes and other eukaryotes, for instance it translates UAA and UAG into tyrosine and not into STOP codons (35). By retaining organelles from its cryptophyte prey, *M. rubrum* can use the prey nucleus to serve the chloroplast gene products using the standard code. Given this is possible for several cryptophytes such as *Teleaulax amphioxeia, T. acuta* and *Geminigera cryophila* (TPG clade), the question remains as to why only these taxa and no other cryptophytes apparently can be exploited (15). *M. rubrum* is known to feed on cryptophyte species belonging to different clades, but cannot utilize them for growth and photosynthesis, with the exception of the TPG clade (15, 36).

The construction of reference gene sets separately for *M. rubrum* and *T. amphioxeia* allowed us to compare the gene compositions of the host and its prey in this endosymbiotic system. By mapping the *M. rubrum* and *T. amphioxeia* genes to KEGG pathways, we found that the majority of pathways were present in both species, whereas *M. rubrum* lacked genes involved in photosynthesis, antenna proteins for photosynthesis and carotenoid biosynthesis (Fig. 6A), although we could not completely rule out the possibility that these genes were presented in the *M. rubrum* genome but not expressed. Interestingly, we also found that the pathways related to the biosynthesis or metabolism of some essential compounds such as lysine, glycan and several kinds of vitamins (pantothenate, riboflavin and biotin) were absent or non-expressed in *M. rubrum* (Fig. 6A), indicating that *M. rubrum* has to obtain the metabolic potential for these compounds from the prey. This explains its dependency on grazing. Of note, most of these *M. rubrum*-absent pathways were actually up-regulated in *T. amphioxeia* after sequestration (Fig. 6A), implicating that *M. rubrum* is able to obtain these nutrients from its cryptophyte prey without digesting the acquired organelles, which is a critical step towards a permanent endosymbiosis. It can be speculated that, in addition to the nucleomorph of the chloroplast, these micronutrients are the reason why *M. rubrum* sequesters exclusively species of the TPG clade.

**Fig. 6.**
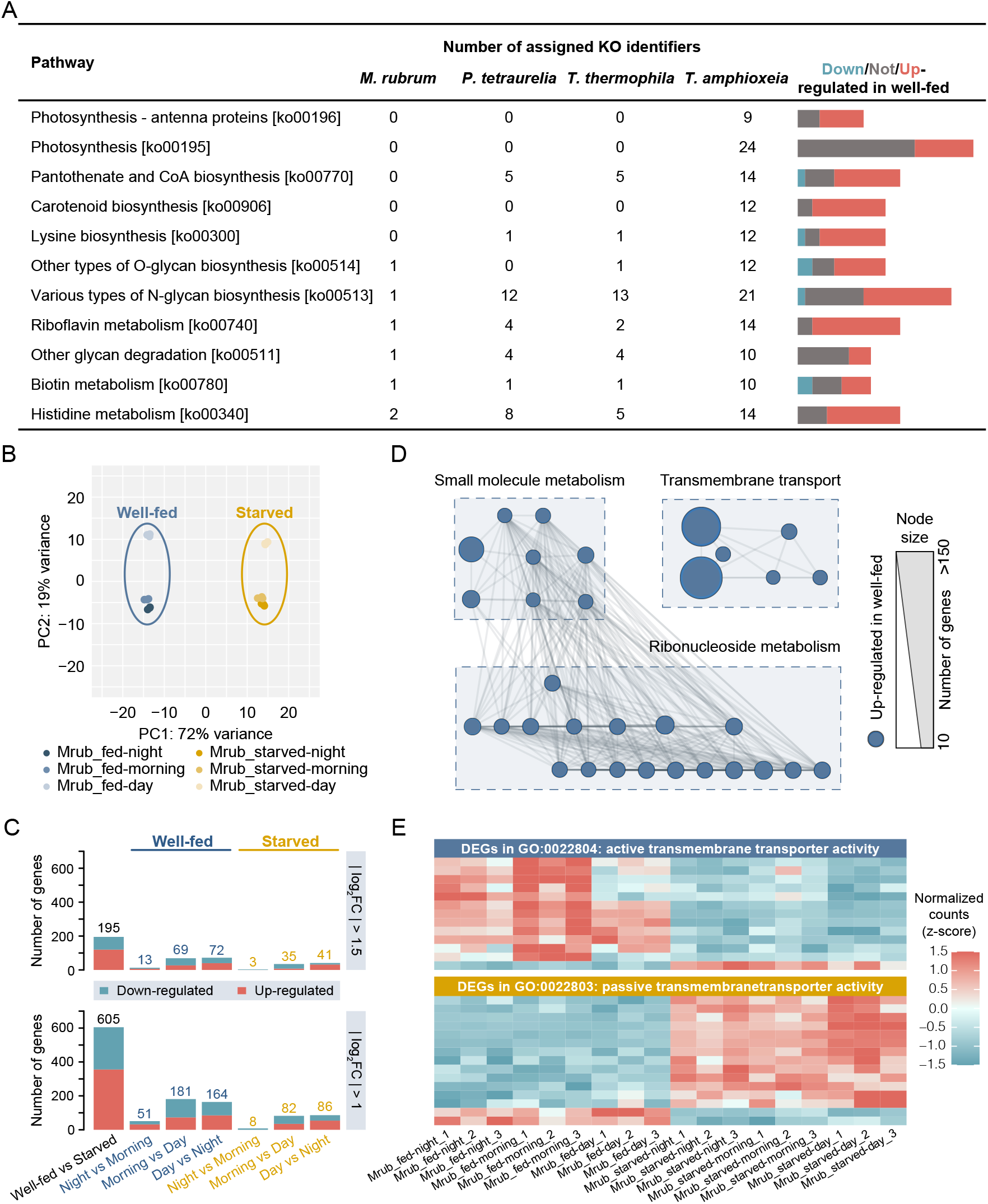
Transcriptional changes in *M. rubrum* upon sequestration in different light conditions. (*A*) comparison of the presence of genes from selected pathways in the ciliates *M. rubrum, Paramecium tetraurelia*, and *Tetrahymena thermophile* and the cryptophyte *T. amphioxeia* with differential expression of *T. amphioxeia* genes upon sequestration (free-swimming versus well-fed), showing services provided by *T. amphioxeia* to *M. rubrum*. (*B*) principal component analysis of *M. rubrum* genes. (*C*) barplot showing the amount of significantly differentially expressed *M. rubrum* genes (|log_2_FC| > 1.5 or |log_2_FC| > 1) according to prey and light conditions. (*D*) GO enrichment analysis of *M. rubrum* genes upregulated in the well-fed samples visualized as an enrichment map. (*E*) heat map showing the differential expression (up/down fold change > 1.5) of active and passive transmembrane transporters in well-fed and starved *M. rubrum* cells.

Next, we investigated the transcriptional changes of *M. rubrum* genes in response to different light conditions and prey availability (well-fed or prey-starved). PCA with the *M. rubrum* gene expression matrix revealed that the *M. rubrum* samples were clustered according to light condition as well as according to prey availability (Fig. 6B). The response to the supply of prey was stronger than the response to light (Fig. 6B). This finding was confirmed by a WGCNA analysis, which uncovered 10 co-expression modules. The two largest modules comprised up to 35% of the input genes and enriched *M. rubrum* genes that were prevailingly down- and up-regulated after starvation (Supplementary Fig. S7). However, DEG analyses using the same cutoff as the *T. amphioxeia* genes (|log_2_FC| > 1.5 and FDR < 0.01), or even lower cutoff (|log_2_FC| > 1 and FDR < 0.01), only identified a small number of genes as DEGs between well-fed and prey-starved *M. rubrum* samples (Fig. 6C). Actually, the negligible number of DEGs identified between samples from different light conditions suggests that *M. rubrum* is not sensitive to light changes, and that the few responses to light take comparatively long time (Fig. 6B, C).

To uncover the functional preference of *M. rubrum* genes in response to prey availability, we conducted functional enrichment analysis for genes clustered in the two largest co-expression modules of the WGCNA analysis. Module 1 comprised 3,285 genes that got downregulated upon prey starvation (Supplementary Fig. S7). Module 2 contained 2,387 *M. rubrum* genes that got upregulated upon starvation (Supplementary Fig. S7). Genes downregulated upon prey starvation (upregulated in well-fed condition) were enriched in small molecule metabolism, ribonucleoside metabolism and transmembrane transport (Fig. 6D). Interestingly, genes related to active transmembrane transporter activity (GO:0022804; adjusted *p* = 0.001) were enriched in module 1 (Supplementary Table 10). This indicates that *M. rubrum-*derived active transmembrane transporters play an important role in well-fed *M. rubrum* (Fig. 6E). In contrast, passive transmembrane transporter activity is more prominent in prey starved *M. rubrum* (Fig. 6E), indicating that well-fed *M. rubrum* is transporting molecules among different cell compartments and actively coordinating biological processes of itself and the cryptophyte prey within the cell.

## Conclusions

We found very strong transcriptional changes of *T. amphioxeia* genes after sequestration by *M. rubrum.* Upregulated prey genes were related to photosynthesis and metabolism, as well as biosynthesis of lysine and glycan, several kinds of vitamins and gene replication. These processes provide a gain for the host, *M. rubrum* and demonstrates its prey dependency. Light dependent transcriptional regulation of *T. amphioxeia* genes found in free-swimming condition got lost upon sequestration. The transcriptional pattern of *T. amphioxeia* genes in well-fed and prey-starved *M. rubrum* was highly similar, indicating that *M. rubrum* can only induce the expression of one particular pattern out of the acquired prey nucleus. *M. rubrum* shows only very few adjustments in its gene expression in response to different light conditions. Noticeable is the upregulation of active transmembrane transporters in well-fed *M. rubrum* and the role of passive transmembrane transporters in starved *M. rubrum.*

## Materials and Methods

### Cultures

Cultures were established from single-cell isolates of *Teleaulax amphioxeia* (SCCAP K-1837, collected in Elsinore Harbor, Denmark), and *Mesodinium rubrum* (MBL-DK2009 collected in September 2009 in Elsinore Harbor, Denmark). Cultures (*T. amphioxeia, M. rubrum* fed *T. amphioxeia*) were kept in triplicates and grown in glass bottles in F/2 medium at 15 °C in a light/dark cycle of 16/8h with a light intensity of 100 μmol photons m−2s−1. During the exponential phase of growth, the ciliates were transferred to new media when cell concentrations reached 5000 ml-1 or more.

### RNA extraction

For RNA extraction, cultures were harvested in full light (7 hours into the light cycle), in darkness (6 hours into dark cycle) and in the transition between dark and light (20 minutes into the light cycle). Cells of *M. rubrum* were harvested in a well-fed and a starved stage.

For the well-fed condition, we checked before extraction that no free cryptophyte cells remained in the medium and that at least 90% of all *M. rubrum* cells contained a cryptophyte nucleus. This was done by staining the nuclei with Hoechst reagent (#33342, Thermo Fisher Scientific, Waltham, USA), and checking 20 stained cells under a fluorescent microscope. Harvesting of starved cells was done approximately four weeks after the last cryptophytes had been seen in the culture. We confirmed the loss of cryptophyte nuclei by staining with Hoechst reagent and checking for prey nuclei under a fluorescence microscope. Cells were harvested after at least 90% of all *M. rubrum* cells had lost their cryptophyte nucleus. Cells were harvested by centrifugation in 10ml glass tubes at 3220 rcf for 10 minutes (see Supplementary Table 11 for cell numbers in each harvest). Pellets were transferred to 1.5 mL LoBind Eppendorf tubes and liquid nitrogen was directly added onto the pellets. The Eppendorf tubes were stored on ice without allowing the pellets to thaw until the lysis buffer was added. RNA was extracted using the column based Exiqon Cell and Plant RNA Isolation Kit (#300110, Exiqon, Vedbæk, Denmark) following the ‘plant’ protocol. In addition, a separate round of harvest has been transferred to hot Trizol and stored at −80 °C as backup. Two samples (10 and 11) from this backup have been used for RNA extraction using the Trizol method. Extracted RNA was stored at −80 °C until library preparation for sequencing.

### DNA extraction

For DNA extraction *T. amphioxeia* cells as well as starved *M. rubrum* (fed *T. acuta)* cells were harvested as described above and DNA extracted using a KingFisher Duo Prime System (#5400110, Thermo Fisher Scientific, Waltham, USA) using the Plant DNA Kit and following the manufacturers recommendations.

### Library construction and sequencing

The RNA-seq libraries were mainly prepared using the MGIEasy RNA Library Prep Set (V1.0, MGI Tech) with 1 ug total RNA as input and sequenced on the BGISEQ-500RS platform using the PE100 chemistry according to the standard protocols provided by MGI Tech Co., Ltd (Shenzhen, China). The only exception was one of the three biological replicates of Tamp-day, of which the amount of total RNA was less than 1 ug and failed to meet the requirement of the MGI kit. The RNA-seq library of this sample was prepared using the TruSeq Stranded mRNA LT Sample Prep kit (RS-122-2101, Illumina) with 500 ng total RNA as input, and sequenced on the Illumina HiSeq 4000 platform using the PE100 chemistry, according to the standard Illumina protocols (San Diego, CA, USA).

The DNA sequencing libraries of free-swimming *T. amphioxeia* and starved *M. rubrum* were prepared using the MGIEasy DNA Library Prep Kit (V1.1, MGI Tech) with 1 μg genomic DNA as input, and sequenced on the BGISEQ-500RS platform using the PE100 chemistry according to the standard protocols provided by MGI Tech Co., Ltd (Shenzhen, China).

### Quality control of raw sequencing data

Prior to subsequent analyses, all the DNA- and RNA-seq raw reads were passed to SOAPnuke (v1.5.3) (37) for quality control by removal of adapter-contaminated reads and low-quality reads. Specifically, all the DNA-seq data were filtered by SOAPnuke with parameters *−n 0.02 −l 20 −q 0.3 −Q 2 −G −d*. For the RNA-seq data, we generated two versions of clean data. The first one was used for de novo transcriptome assembly and generated by parameters *−n 0.02 −l 20 −q 0.3 −p 1 −t 15,0,15,0 −Q 2 −G −d* (i.e. removing adapter-contaminated, low-quality and duplicated reads). The second version was used for gene expression measurement and generated by parameters *−n 0.02 −l 20 −q 0.3 −p 1 −t 15,0,15,0 −Q 2 −G* (i.e. removing adapter-contaminated and low-quality reads but keeping duplicated reads).

### Construction of reference gene sets for *T. amphioxeia* and *M. rubrum*

A hierarchical strategy was employed to construct the reference gene sets for *T. amphioxeia* and *M. rubrum* with the RNA-seq and DNA-seq clean data.

#### (i) De novo transcriptome assembly for each RNA sample

The clean RNA reads (without duplicates) of each sample were first assembled using Trinity (v2.4.0) (38, 39) with parameters *--min_contig_length 150 --min_kmer_cov 2 -- min_glue 3*. Then the highly similar sequences were clustered, and the redundant transcripts were removed from each transcriptome assembly using cd-hit-est (v4.6.8) (40, 41) with a sequence identity threshold of 0.95. Finally, the clean RNA reads (with duplicates) were aligned to each assembly to quantify the abundance of each gene defined by Trinity (i.e. a transcript cluster) using Salmon (v0.13.1) (42) with parameters *--validateMappings −l IU –allowDovetail*. The lowly expressed genes with TPM < 1 were removed from each transcriptome assembly, then only the longest transcript of each gene was kept. Detailed statistics for the transcriptome assembly for each RNA sample is available in Supplementary Table 2.

#### (ii) Assignment of transcripts to *T. amphioxeia* and *M. rubrum*

Species assignment was conducted based on the average nucleotide identity (ANI) of each transcript in relative to the DNA sequences of *T. amphioxeia* and *M. rubrum* estimated by Mash (v2.1) (43, 44). To eliminate potential prokaryotic contamination in the transcriptomes, we also built a data set containing 10,243,458 prokaryotic nucleotide sequences by extracting all Archaea and Bacteria sequences from the NCBI nt database (release 20190315). Specifically, we sketched at most 5,000 non-redundant 21-mers from each transcript, and compared them with the non-redundant 21-mers generated from all of the *T. amphioxeia* DNA reads, *M. rubrum* DNA reads and the prokaryotic nucleotide sequences, respectively, to estimate ANI of *T. amphioxeia* (ANItamp), *M. rubrum* (ANImrub) and prokaryotic (ANIprok) for each transcript. Of note, even so, more than 90% of the cells in the starved *M. rubrum* cultures had lost the cryptophyte nucleus, genomic sequences of *T. amphioxeia* was still detectable in the starved *M. rubrum* DNA reads (although occurring at low abundances). Considering this, a transcript was assigned to: (1) *T. amphioxeia* if ANItamp ≥ 0.95 and ANIprok < 0.95; (2*) M. rubrum* if ANItamp < 0.95, ANImrub ≥ 0.95 and ANIprok < 0.95; (3) unknown sequences for other conditions (Supplementary Table 2). The unknown sequences were discarded from subsequent analyses.

#### (iii) Hierarchical removal of redundant transcripts

The reference gene sets of *T. amphioxeia* and *M. rubrum* were generated by combining *T. amphioxeia* and *M. rubrum* transcripts from all the samples, respectively. The highly similar sequences were clustered, and the redundant transcripts for the same species were removed using cd-hit-est (v4.6.8) (40, 41) with a sequence identity threshold of 0.95. To further remove redundant transcripts that failed to be clustered by cd-hit-est (caused by alternative splicing such as exon skipping, intron retention, etc.), two rounds of Minimap2 spliced alignment and MCL clustering processes were performed. Specifically, pairwise spliced alignment for all transcripts was generated using the all-vs-all mode of Minimap2 (v2.10-r764) (45) with parameters *−aX −x splice*. Then, a graph was built with the best alignment of each query that satisfied identity > 0.95 and coverage > 70% against the shorter sequence. The vertices of the graph were the transcripts and the edges were weighted by identity x coverage. Next, the graph was inputted to Markov Clustering algorithm (mcl v14-137) (46) to cluster similar sequences with the default power and inflation setting, and the longest transcript of each cluster was kept as the representative. The Minimap-MCL process ran iteratively based on the cluster representatives generated by the last iteration until no more pairwise alignments satisfying the threshold were found. Then a second round of the Minimap-MCL process was performed with the threshold identity > 0.98 and coverage of the shorter sequence > 50% to generate the final non-redundant *T. amphioxeia* and *M. rubrum* reference gene sets. The longest ORF for each gene was detected by TransDecoder (v5.5.0) (39) with parameters *-m 50 --genetic_code universal* for the *T. amphioxeia* genes and *-m 50 --genetic_code Mesodinium* for the *M. rubrum* genes.

#### (iv) Estimation of the representativeness of the reference gene sets

To evaluate the representativeness of the reference gene sets, we aligned the RNA clean reads (with duplicates) from each of the *T. amphioxeia* samples to the *T. amphioxeia* genes, and aligned the reads from each of the *M. rubrum* samples to the collection of *T. amphioxeia* and *M. rubrum* genes using BWA-MEM (v0.7.16) (47) with default parameters. Then, the numbers and ratios of reads being aligned, being uniquely aligned (as defined by mapping quality ≥ 30), and being aligned in proper pairs, in relative to the total numbers of inputted reads, were counted by samtools flagstat (SAMtools v1.7) (48, 49). The completeness of the gene sets was assessed by the Benchmarking Universal Single-Copy Orthologs (BUSCO v3) (50) with the eukaryota odb9 database (Supplementary Table 12).

### Identification of *T. amphioxeia* genes present in prey-starved *M. rubrum* DNA data

To determine whether the whole prey nuclei were kept inside *M. rubrum* (Fig. 3A), DNA read alignment and alignment-free k-mer screening methods were used to identify *T. amphioxeia* genes in the two starved *M. rubrum* DNA samples. For the alignment based method, we aligned the DNA reads of the two starved *M. rubrum* DNA samples to the *T. amphioxeia* gene set using BWA-MEM (v0.7.16) (47) and calculate the coverage of each gene using genomeCoverageBed (v2.26.0) (51) with default parameters. *T. amphioxeia* genes with coverage larger than 70% were considered present in the starved *M. rubrum* cells. The alignment-free k-mer screening method was described in the above section “Assignment of transcripts to *T. amphioxeia* and *M. rubrum*”. The *T. amphioxeia* genes present in the starved *M. rubrum* DNA samples were defined as ANItamp ≥ 0.95 and ANIprok < 0.95.

### Expression level quantification

We aligned the RNA clean reads of each sample to the database containing all *T. amphioxeia* and *M. rubrum* genes and quantified the abundance of each gene using Salmon (v0.13.1) (42) with parameters *--validateMappings −l IU --allowDovetail*. Of note, more than 99.98% of the aligned reads from the free-swimming *T. amphioxeia* samples were mapped to the *T. amphioxeia* genes by Salmon, highlighting the reliability of our species assignment process described above. To eliminate the big disparity in the data amount originated from *T. amphioxeia* across different sample groups (Fig. 3D) for subsequent analyses, we downsampled the clean data of each sample to adjust the number of *T. amphioxeia* aligned reads in each sample to at most ~40M. The raw counts of the *T. amphioxeia* genes in all the 27 samples were collected (Supplementary Table 4). Then, we balanced the data amount originated from *M. rubrum* across all *M. rubrum* samples in the same way and generated the raw counts of the *M. rubrum* genes in all the 18 *M. rubrum* samples (Supplementary Table 4).

### Transcriptome features of *M. rubrum*

With the conclusion that *M. rubrum* keeps all the genetic material from the prey (Fig. 3A), we further investigated how actively the *T. amphioxeia* genes were transcribed in the *M. rubrum* samples (Fig. 3B) and the component features of the global *M. rubrum* transcriptomes (Fig. 3C and 3D). Specifically, three matrices were first collected from the Salmon quantification output: EffectiveLength of each *T. amphioxeia* gene in each sample, Counts of each *T. amphioxeia* gene in each sample and TPM of each *T. amphioxeia* and *M. rubrum* gene in each *M. rubrum* sample. Next, the *T. amphioxeia* TPM matrix were then re-calculated with the EffectiveLength and Counts matrices according to formula: TPM_i_ = 10^6 * (Counts_i_ / EffectiveLength_i_) / [Σ (Counts_i_ / EffectiveLength_i_)]. The genes with TMP ≥ 1 were deemed to be actively transcribed. Finally, we calculated the proportion of actively transcribed *T. amphioxeia* genes in all samples with the re-calculated *T. amphioxeia* TPM matrix, the ratio of actively transcribed *T. amphioxeia* and *M. rubrum* gene numbers (NumGenes_*Tamp*_/(NumGenes_*Tamp*_+ NumGenes_*Mrub*_) vs. NumGenes_*Mrub*_/(NumGenes_*Tamp*_+ NumGenes_*Mrub*_)) and accumulated the abundance of actively transcribed *T. amphioxeia* and *M. rubrum* genes (Σ TPM_*Tamp*_ / (Σ TPM_*Tamp*_+ Σ TPM_*Mrub*_) vs. Σ TPM_*Mrub*_ / (Σ TPM_*Tamp*_+ Σ TPM_*Mrub*_)) in each *M. rubrum* sample with the *T. amphioxeia* and *M. rubrum* TPM matrix. The proportion of actively transcribed *M. rubrum* genes at different prey availabilities (Fig. S1) was counted in the same manner as Fig. 3B.

### Identification of differentially expressed genes (DEGs)

The DEG candidates between sample groups were detected by DESeq2 (v1.22.0) (52). First, the library sizes across samples were normalized using the default median ratio method, taking the raw counts as input. Next, to obtain dispersion estimates, the type of fitting of dispersions to the mean intensity was set to be parametric. Then, we used Wald significance tests (nbinomWaldTest) for model fitting and test statistics. The Benjamini-Hochberg False Discovery Rate (FDR) was employed to adjust p-value for multiple test correction. Finally, the significant DEGs were defined with the criteria of basemean ≥ 20, adjusted p-value < 0.01 and |log_2_ (fold change)| > 1.5 (|log_2_ (fold change)| > 1.5 and |log_2_ (fold change)| > 1 were both used to detect significant *M. rubrum* DEGs).

### Principal component analysis (PCA) clustering

We used the normalized counts as described in the above DEG section to perform PCA clustering. We filtered the gene counts with narrow variance (standard deviation of normalized count < 10 across all samples) and then vst (variance stabilizing transformation provided by DESeq2) transformed the normalized counts with parameter *blind=FALSE*. Finally, we generated the PCA plots with the vst transformed matrices using plotPCA function provided by DESeq2.

### Identification of co-expression modules

Co-expression network analyses were performed using the weighted gene correlation network analyses R package (WGCNA v1.68) (53). After filtering lowly expressed genes (mean normalized count < 10 across all samples), the normalized read count matrices were vst transformed with parameter *blind=FALSE*. Then the vst transformed matrice were passed to the blockwiseModules function implemented in the WGCNA package for identification of the signed co-expression modules with parameters *maxBlockSize = 50000, networkType = signed, minModuleSize = 100, minKMEtoStay = 0.6, minCoreKME = 0.5, mergeCutHeight = 0.15, numericLabels = TRUE, pamRespectsDendro = FALSE* and *power = 16* for the *T. amphioxeia* genes and *power = 18* for the *M. rubrum* genes.

### Functional annotation and enrichment analysis

We aligned the gene sequences to the UniProtKB/Swiss-Prot database (release 20190408) with BLASTX (blast-2.2.26) using parameters *-F F -e 1e-5*. The best hit of each query was retained based on the BLASTX bit score. The GO annotation of the best aligned UniProt protein was then assigned to the query gene. To determine what pathways the genes might be involved in, gene sequences were searched against the KEGG database (v89.1)(54) with BLASTX (blast−2.2.26) using parameters *−F F −e 1e-5*. The best hit of each query was retained based on the BLASTX bit score.

A set of better quality and reliable functional annotations was generated (the BLASTX hits with e-value lower than 1e-10) for the functional enrichment analyses. Hypergeometric tests were employed to examine whether a list of DEGs was enriched in a specific GO term in relation to background genes as previously described(55), by comparing the number of target DEGs annotated to this GO term, the number of target DEGs not annotated to this GO term, the number of background genes (i.e. all the *T. amphioxeia* or *M. rubrum* genes excluding the target DEGs) annotated to this GO term, and the number of background genes not annotated to this GO term. P-values were adjusted for multiple testing by applying FDR (56), and enriched GO terms were considered significant for adjusted p-value < 0.05. The GO enrichment results were visualized with EnrichmentMap (v3.2.1) (57) in Cytoscape (v3.7.2) (58).

KEGG enrichment analysis was done using the same principle as the GO enrichment. The regulation of gene expression involved in the enriched KEGG pathways was visualized Pathview (v1.23.3)(59).

### Comparison of the presence of genes involved in the pathways in the ciliates and the cryptophyte

To compare the biological process compositions of the host *M. rubrum* and its prey *T. amphioxeia* in this endosymbiotic system, we first extracted and counted the KO (KEGG Ontology) terms and pathways present in the reference ciliates *Paramecium tetraurelia* and *Tetrahymena thermophile* directly from the KEGG database (v89.1). For the host *M. rubrum* and its prey *T. amphioxeia*, we aligned the gene sequences to the KEGG database (v89.1) with BLASTX (blast-2.2.26) using three sets of parameters: (a) *−F F −e 1e-5*, (b) *−F F −e 1e-10* and (c) *−F F −e 1e-20*. The KO terms associated with the pathways of the best aligned KEGG protein were then assigned to the query genes. The pathways involving less than 5 genes with BLASTX hits (e-value < 1e-5) in *M. rubrum* and at least 20 genes with BLASTX hits (e-value < 1e-20) in *T. amphioxeia* were considered absent or non-expressed in the host *M. rubrum* and provided by the prey *T. amphioxeia* in the endosymbiotic system. The number of present KO terms in the selected pathways in *M. rubrum* and *T. amphioxeia* shown in Fig. 6A were counted with the BLASTX hits (e-value < 1e-10).

## Data availability

The raw sequencing reads produced in this study are deposited in the CNGB Nucleotide Sequence Archive (CNSA) with accession number CNP0000925 (https://db.cngb.org/cnsa/). The nucleotide sequences and functional annotations of the reference gene sets for *Teleaulax amphioxeia* and *Mesodinium rubrum* are deposited in the figshare repository.

## Acknowledgements

The Danish Council for Independent Research funded this project (grant number 4181-00484 to PJH). QL acknowledges the Major scientific and technological projects of Hainan Province (ZDKJ2019011). LGC was supported by the Carlsberg Foundation (2012_01_0515).

## Author contributions

PJH, NL, UJ and AA conceived the work. AA, KD, MK, LGC did the culturing; AA and LGC extracted RNA; QL, YZ and XZ did the sequencing; HC, AA and QL analyzed the data under the supervision of SL and NL; AA, HC and QL drafted the manuscript. All authors contributed to writing of the final version of the manuscript.

## Conflict of Interest

The authors declare no conflict of interest.

## Notes

### Competing Interest Statement

The authors have declared no competing interest.

